# Physics-Informed Neural Network Methods for Predicting Plant Height Development

**DOI:** 10.64898/2026.01.14.699475

**Authors:** Yingjie Shao, Fred van Eeuwijk, Carel F.W. Peeters, Olivia Zumsteg, Ioannis N. Athanasiadis, George van Voorn

## Abstract

Plant growth is a dynamic process affected by genes and growing environment, with all kinds of interactions between them. These complex relationships make the prediction of plant growth challenging. We propose a hybrid modelling framework that combines a logistic ordinary differential equation model with a Long Short-Term Memory (LSTM) neural network, resulting in a Physics Informed Neural Network (PINN). While PINNs have been widely applied to physical dynamical systems, their use in modelling the dynamics of plant growth systems is still largely unexplored. We illustrate the construction of a PINN on plant height data in wheat and compare its performance with alternative models for longitudinal plant data. All temporal prediction models only require time and temperature as input. Among a set of competing models, our PINN had the lowest average root mean squared error (RMSE) of prediction and the smallest standard deviation across multiple random initialisations. Therefore, we conclude that incorporating biological growth constraints into data-driven growth models can enhance prediction accuracy of longitudinal plant traits.

**Highlights:**

- Integrating plant growth equations into a temporal neural network improves plant height growth prediction over ordinary differential equations and machine learning models, especially when training data are limited.

## 2. Introduction

Predicting how existing or new crop genotypes will grow under novel environmental conditions is one of the critical challenges in agriculture. Accurate prediction is important for optimising crop growth under diverse conditions, supporting crop breeding, crop management decisions, and other applications (Xiong et al., 2022). Plant growth results from multiple genetic and environmental interactions that can be non-linear and time-dependent (Napier et al., 2023), which increases the complexity of prediction. The increasing availability of high-throughput phenotyping (HTP) data allows modellers to train temporal prediction models for such tasks (Ranđelović et al., 2023; Roth et al., 2023). In particular, the non-destructive, sensor-based longitudinal data from HTP platforms provide continuous monitoring of plant traits alongside environmental covariates, offering rich information for temporal prediction. Building on these opportunities, this work aims to develop and compare approaches for plant growth prediction.

Data-driven and process-based approaches are the two main strategies for plant growth prediction, and both have advantages and limitations. Although classical statistical methods have been used for crop phenotype prediction, they are generally less flexible than neural networks and are not considered in this study. Data-driven neural network models can learn complex patterns from HTP data, but they do not necessarily incorporate or obey physical mechanisms (Wu et al., 2024). The lack of mechanistic descriptions of crop growth can limit the ability of these models to extrapolate predictions beyond the data domain. Such extrapolations include predicting plant development further within the growing season, as well as predicting how specific genotypes grow in a new season or different environment, or how newly developed genotypes may perform. Conversely, process-based models encode mechanistic knowledge explicitly, but their fixed model structures can reduce flexibility and limit their ability to capture complex patterns present in real-world data (Droutsas et al., 2022).

Physics-Informed Neural Network (PINN) models present a promising route to address these limitations and combine the advantages of data-driven and mechanistic approaches by incorporating domain knowledge into neural network models. Neural networks are data-driven models that have become more popular with the increasing availability of HTP data (Danilevicz et al., 2022). HTP data allow the neural network models to ‘learn’ correlations from datasets for temporal prediction by identifying implicit functions between inputs (environmental factors, etc.) and outputs (plant traits). Neural network structures specifically designed for temporal analysis, such as Recurrent Neural Networks (RNNs) and Long Short-Term Memory (LSTM) architectures, process temporal data and utilise information from previous time steps for current time-step prediction (Hochreiter and Schmidhuber, 1997; Marhon et al., 2013). LSTM-based models have been used to capture temporal dependencies in various applications in agriculture (Khaki et al., 2020; Siami-Namini et al., 2019; Wang et al., 2025). In this work, we integrate the temporal modelling capabilities of LSTMs with explicit mechanistic knowledge using a Physics-Informed Neural Network (PINN) framework.

In a PINN structure, domain knowledge is encoded as explicit relationships, typically using ordinary differential equations (ODEs). Ordinary differential equations relate the rates of change in variables, such as plant traits, to inputs such as environmental factors and observational plant traits. This makes ODE-based equations suitable for the description of time-dynamic and non-linear processes (Daun et al., 2008), such as those that underlie plant growth. In addition, ODEs are smoothly differentiable, which is convenient for the backward propagation needed for training neural network models. PINN models use ODEs in the loss function, where the ODEs represent relevant dynamic processes (Nathasarma and Roy, 2023; Raissi et al., 2017). The ODE component limits the solution space, which is especially helpful with limited or irregularly sampled training data (Chen et al., 2023; Cuomo et al., 2022; Raissi et al., 2019). However, any ODE will not necessarily cover all relevant processes; the data-driven part of the PINN model is therefore intended to augment missing information (Jhutty and Hernandez-Vargas, 2022).

PINNs have been successfully applied to biological systems, such as modelling soil microbiota growth (Cuomo et al., 2025) and simulation of soil thermal dynamics without input of soil thermal properties (Xie et al., 2024). Although used for dynamic systems, most reported PINNs are built with feed-forward architectures (Cuomo et al., 2022), which treat time as an explicit input and ignore sequential temporal structure, limiting their ability to capture temporal dependencies in dynamic biological processes. Furthermore, PINNs have not yet been studied in agricultural contexts for modelling dynamic plant traits. This represents a missed opportunity given the increasing availability of time-resolved high-throughput phenotyping data.

In this paper, we construct PINN-based models to evaluate their ability to forecast plant traits under future environmental conditions based on crop trial data. We compare our PINN models with two ODE-based models, a classical machine learning (ML) model type, and a time-explicit neural network type. The evaluation is based on prediction accuracy, robustness, and the potential to extend the model to forecast the growth of existing or new genotypes in new environments. The data used to train the models are described in Section 3.1. The models are described in Section 3.2. The comparison metrics are given in Section 3.3. The main model comparison results are given in Section 4. The results are discussed and linked to current limitations and potential future studies in Section 5.

## 3. Material and Methods

To develop a PINN model for predicting plant height of multiple genotypes under future temperature conditions, this section follows the logical sequence of the model prediction process, from input to output. We propose our PINN model: an LSTM-based Logistic ODE-informed PINN model (*Logi-PINN*), which has not been explored in dynamic plant growth prediction tasks. The proposed *Logi-PINN* was evaluated by model comparison with two different setups: single-genotype models (one model for each genotype separately) and multiple-genotype models (one model for multiple genotypes simultaneously).

We begin this section by describing the HTP dataset used in this study and the pre-processing steps applied. The genotype encoding section explains how we included genotype information in our multiple-genotype model. In the Model section (Section 3.2), we introduce the five model structures, including our proposed *Logi-PINN*, with detailed descriptions of key components (such as LSTM units) provided in the supplementary data (Figure S1.1). Finally, we describe the evaluation metrics used to assess prediction performance and the impact of the physics loss components. To facilitate understanding of the equations presented in the following section, we provide a notation table for easy reference in Table S2.

### 3.1. Dataset

#### 3.1.1. Data description

Plant height results from interactions between genetic factors and time-dynamic environmental factors, such as temperature, irradiance, and water content (Amalova et al., 2024; Miao et al., 2024). Plant height reflects plant development and depends on a limited number of genes (Wu et al., 2010). It indirectly contributes to yield by affecting photosynthesis through leaf area index and biomass, which have been used for yield prediction (Gracia-Romero et al., 2023; Tao et al., 2020). Moreover, accurately predicting plant height growth also helps in precise management (Jayakumari and Nidamanuri, 2024), such as irrigation and fertilisation, to maintain height within an optimal range. Plant height can be automatically measured with high-throughput phenotyping (HTP) technology, resulting in high-temporal-resolution data that can be used for constructing prediction models.

We trained and evaluated our models on a subset of a real-world dataset of wheat growth collected by ETH Zürich on a field phenotyping platform (FIP) from 2018 to 2022 (Roth et al., 2025). The HTP dataset of the field experiment consists of frequent but irregular measurements of phenotypic variables across the growing season and regularly measured environmental variables, and different variables often do not have the same temporal resolution (Geng et al., 2024; Roth et al., 2021). This dataset covers hundreds of genotypes and includes hourly measurements of air temperature measured two metres above ground, genetic markers, and estimated longitudinal plant height (Roth et al., 2025). The dataset has been used for modelling growth responses to temperature (Roth et al., 2022), and shows the relationship between elongation and temperature for different genotypes (Roth et al., 2024). Plant height in the dataset was extracted from RGB images captured by drones, resulting in approximately 40 plant height data points per plot during the growing season. Temperature was selected as input for our temporal prediction model, as it is an important environmental factor that affects enzyme-controlled reactions within plants, such as photosynthesis, leading to changes in plant height (Kronenberg et al., 2021; Proctor, 1982). Typically, two replicates per genotype were grown each year and assigned to plots following an experimental design; however, some genotypes included additional replicates, as indicated in Figure S5.1. All replicates were treated as independent samples during model training to increase the number of target labels, while replicate measurements corresponding to the same genotype and year were averaged for model evaluation, as they were grown under identical temperature conditions.

#### 3.1.2. Data pre-processing

The four-year plant height time series data, spanning from October to July, were aligned based on sowing date and kept at daily intervals by filling gaps with *NA*s between measured dates, resulting in 285-day time series. Because the data concern winter wheat, the first 115 days (up to around March) were then removed to exclude non-informative records before air temperature starts to increase and elongation begins (Figure S3.2), resulting in *n*_*t*_(= 170)-day time series with approximately 30–40 non-*NA* values per plot per year. Temperature data consisted of daily averages to align with the plant height observations. To handle missing early-season plant height values introduced by time series alignment and to improve the model’s ability to learn initial conditions, we filled time points preceding the first recorded height in each year with the sequence’s minimum height value. The pre-processing pipeline is illustrated in Figure 1. The processed data were then formatted as model inputs, as described in Section 3.2.

**Figure 1:**
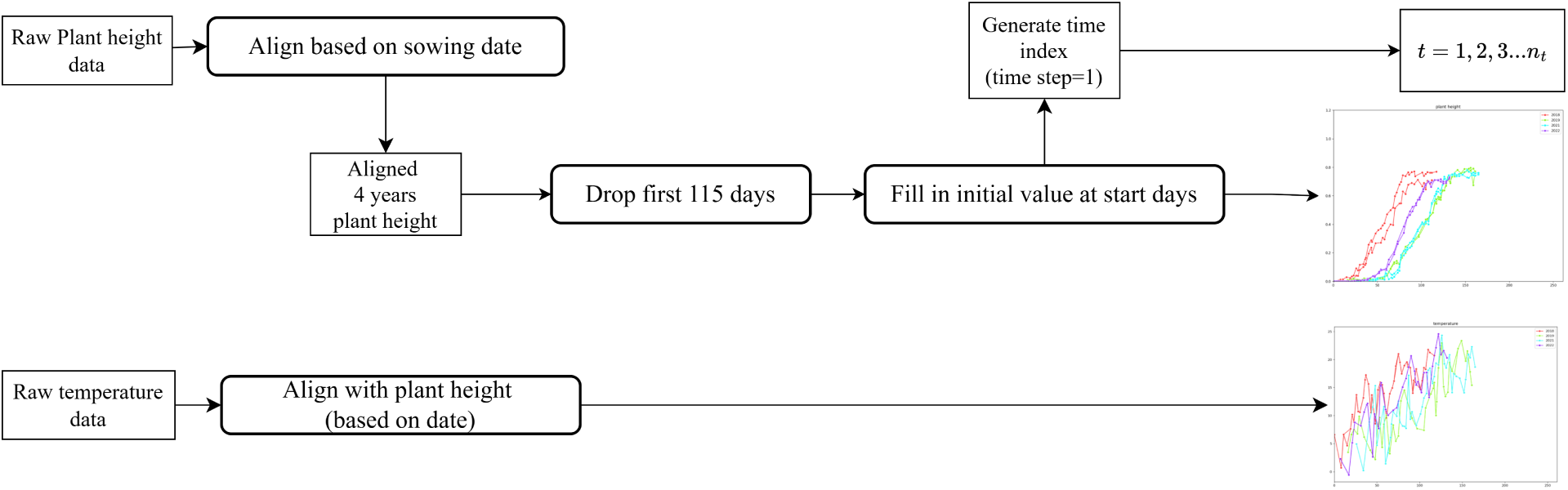
Data Pre-processing. This plot visualises the pipeline from raw temperature and plant height sequences to the processed inputs for model training. The temperature time series was aligned with the plant height series and also kept at the same time resolution. An example of a processed data plot is shown at the end of the pipeline.

#### 3.1.3. Genetic encoding

We selected GABI wheat genotypes (Kollers et al., 2013) that were present in all four years after pre-processing, resulting in a total of *n*_*g*_= 19 genotypes. The genetic information for the dataset is composed of single-nucleotide polymorphism (SNP) marker scores. The genotypes were encoded with two approaches to make genotype-specific predictions: one-hot encoding and kinship matrix encoding. Encoded genotype vectors were used in the model to associate differences in plant height curves with their genotype and also link the genotypes with parameters in the ODE, enabling genotype-specific plant height predictions with the same model. For genotype one-hot encoding, each genotype was converted into a unique length *n*_*g*_ vector containing 0s and 1s, and all genotype vectors were then concatenated, resulting in a sparse matrix in which the similarity between any pair of genotypes after encoding is identical. For kinship matrix encoding, a length *n*_*g*_ vector was used to represent the genomic distance between one genotype and the other genotypes in the dataset. Specifically, we calculated it using the “astle” method from the statgenGWAS package to get a kinship matrix based on marker information for each genotype (Rossum and Kruijer, 2024). The “astle” method is the default for the kinship matrix, which was calculated based on covariance between the scaled SNP scores (William, Astle and David, J. Balding, 2009).

#### 3.1.4. Data split

The dataset was divided into three subsets: two training years, one validation year, and one test year. The training dataset was used to train our models. The validation data was used to select hyperparameters (parameters that are set and not learned from data) and the number of epochs (i.e., the number of iterations before training is stopped). The test dataset was used as an independent dataset to evaluate model performance.

We repeated the data-splitting process six times, covering all possible combinations of training years, with the same 2:1:1 ratio for the training, validation, and test sets. We focused on evaluating model performance when the model is trained using the years 2018 and 2019, which are contrasting in terms of temperature and thus increase diversity in the training data. The justification behind this was to increase data diversification and avoid high similarity in training data (Gong et al., 2018), which was expected to improve the ability to successfully extrapolate to unseen conditions (an example of different years’ plant height can be found in Figure S3.1).

### 3.2. Models

Before introducing the individual model structures, we defined the model inputs and notation used throughout this section. Let **Y** ∈ ℝ^*n_p_*×*n_t_*^ denote the plant height matrix, where *n_p_* is the number of samples (plots) used in a given model and scenario indexed by *p*, and *n_t_* = 170 is the number of time points after pre-processing. The matrix **T** ∈ ℝ^*n_p_*×*n_t_*^ and **J** ∈ ℝ^*n_p_*×*n_t_*^ were obtained by repeating the time-indexed temperature and time vectors across samples. The matrix **T** ∈ ℝ^*n_p_*×*n_t_*^ describes the daily averaged temperatures. Matrix **J** ∈ ℝ^*n_p_*×*n_t_*^ describes the time index (1…170), which is a necessary component for the *Logi-PINN* to calculate physics loss (Equation 5). Genotype information for multiple-genotype models was provided either as a one-hot encoded genotype matrix **G**_OH_ ∈ ℝ^*n_g_*×*n_g_*^ or as a kinship matrix **G**_K_ ∈ ℝ^*n_g_*×*n_g_*^ (*n_g_* = 19). Both representations were used as inputs to the genetic embedding module.

We compared the proposed *Logi-PINN* with four baseline models representing different modelling paradigms: two process-based ODE models, a Logistic ODE (*Logi-ODE*) and a temperature-informed Logistic ODE (*Temp-ODE*); a classical machine-learning model, namely a random forest (*RF*) model; and a data-driven neural network model (*LSTM-NN*). The ODE models served as process-based baselines, with *Logi-ODE* capturing the general growth trend without explicit environmental forcing and *Temp-ODE* incorporating temperature as a direct modifier of growth rate. The *RF* model provided a commonly used non-temporal machine-learning reference in agricultural prediction tasks, while *LSTM-NN* served as a purely data-driven temporal model for assessing the added value of physics-based constraints in *Logi-PINN*.

Models were first evaluated under a single-genotype prediction setting, where separate models were trained for each genotype. Based on these results, multiple-genotype models were subsequently constructed for the two best-performing neural network structures, *LSTM-NN* and *Logi-PINN*, using explicit genetic encoding with all genotype data.

#### 3.2.1. ODE-based plant growth model

As dynamic process-based models, we selected a *Logi-ODE* and a *Temp-ODE*, both proposed by Van Voorn et al. (2023) for modelling biomass. The reason for choosing the *Logi-ODE* is that previous research shows that (modified) logistic growth is suitable for approximating the general plant height dynamic growth patterns (Van Voorn et al., 2023; Jiang et al., 2020). We therefore assumed that the *Logi-ODE* describes the general trend of plant height without considering environmental factors during the growing season. The *Temp-ODE* added a smooth (i.e., continuously differentiable) temperature response curve that modified the growth rate parameter based on temperature input. Note, that this temperature response curve introduced a direct link between ambient temperature and plant height and not between temperature and photosynthesis. The reason was that no photosynthesis data were available. Furthermore, the temperature response curve may differ according to genotype. The *Logi-ODE* function is given as:

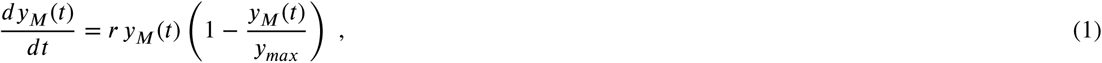

where *y*_*M*_(*t*) is the plant height (m), *r* is the intrinsic growth rate (m/day), and *y*_*max*_ is the maximally attainable height of the plant (m). We randomly initialise *r* between 0.1 and 0.2, and *y*_*max*_ between 0.7 and 0.8 for the *Logi-ODE*, as broader initial ranges often caused convergence issues during the ODE parameter optimisation.

Although the pre-processed dataset assigned small non-zero values to early time points to handle missing observations (Section 2.1.2), these values were *not* used as initial conditions for ODE integration. Instead, the initial condition for the ODE solver was fixed as a small constant, *y*(0) = 0.0001, for all genotypes and years. This choice was made solely for numerical stability, as *y*(0) = 0 is a steady state of the logistic equation and would prevent growth during numerical integration. Biologically, *y*(0) represents an abstract initial height at the start of the modelling window rather than a measured trait value. Because early observed plant height values were likely affected by measurement noise and do not reliably reflect genetic variation, we did not treat *y*(0) as genotype-specific.

We first optimised the ODE parameters *r* and *y*_*max*_ for individual genotypes and years separately. The averaged parameter values from the two training years were then used to predict plant height trajectories for the same genotype in future years. While this approach did not fully exploit the flexibility of process-based models, it provided a consistent and interpretable baseline for comparison with data-driven and hybrid prediction models. Furthermore, the *Logi-ODE* equation was also used as a physics constraint in the proposed *Logi-PINN*.

The *Temp-ODE* is :

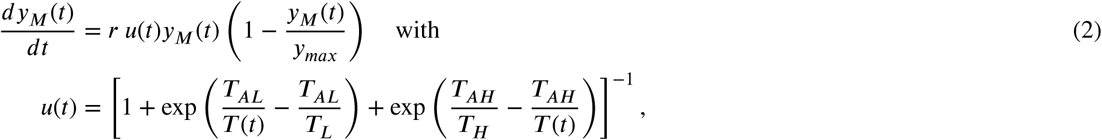

where *T*_*L*_,*T*_*H*_, *T_AL_* and *T_AH_* control the shape of the temperature response curve (*u*(*t*)). This response curve models the effect of temperature on stem elongation by defining a range between lower and upper cardinal temperatures, outside of which growth rate will decrease to zero. *T_L_* and *T_H_* represent the lower and upper boundaries of plant tolerant temperature in Kelvin, respectively, while *T_AL_* and *T_AH_* describe the changing rate of *u*(*t*) when temperatures move towards two boundaries in Kelvin; one can vary the values of these parameters to vary the shape of the response curve to mimic measured response curves (Van Voorn et al., 2023).

Although several *Temp-ODE* parameters vary between genotypes, the model did not include parameters that can be directly linked to SNP information; therefore, it was evaluated only in the single-genotype plant height prediction task. Specifically, the temperature response parameters *T_AL_* and *T_AH_* were fixed and shared across all genotypes and years, while *r*, *y_max_*, *T_L_*, and *T_H_* were fitted separately for each genotype using two-year training data based on their temperature inputs. This reflected the assumption that different genotypes exhibit distinct genetic effects, resulting in different parameter values. The fitted parameter sets were then used to predict plant height trajectories in the validation and test years, making the *Temp-ODE* results comparable with those of other machine-learning prediction models. We used the same parameter initialisation as in the *Logi-ODE* model, with *T_L_* and *T_H_* initialised at 292 and 303 K, respectively, with small random perturbations, while *T_AL_* and *T_AH_* were fixed at 2000 and 60,000 K.

We implemented the code with Python ‘SciPy’ (Virtanen et al., 2020) library. We fitted ODE parameters with the ‘Nelder-Mead’ method, which is a direct search method that optimises parameters by minimising an objective function (root mean squared error) between data and ODE predicted value (Nelder and Mead, 1965). The prediction from ODE was calculated by ODE integration with the ‘LSODA’ method, a numerical ODE solver that is a flexible and reliable method for ODE integration (Petzold, 1983).

#### 3.2.2. RF model

We fitted the *RF* model separately for each genotype. *RF* was selected as it is a widely used ML method in crop prediction, which has given accurate prediction in both regression and classification problems (Dhillon et al., 2023; Jui et al., 2022; Toda et al., 2024). It averages results from multiple decision trees to reduce over-fitting (Ho, 1998) and is relatively insensitive to moderate changes in hyperparameter settings. Because each feature is treated independently for prediction, it does not explicitly use time dependency in input data (Regier et al., 2023). In this way, *RF* represented a minimal performance reference: any alternative model should be able to outperform the relevant *RF* model in terms of prediction ability. The input was treated differently in the *RF* model compared to neural network models. Daily temperature and time index sequences were used as independent features to train the *RF* model. While hyperparameter tuning and tree pruning are often important considerations in *RF* models, we tested several hyperparameter combinations and observed no significant improvement in prediction accuracy. Consequently, we retained the default settings: using ‘squared_error’ as the criterion for feature selection, 100 trees, consideration of all features for the best split, and no additional pre-pruning or post-pruning strategies. Further implementation details can be found in the ‘RandomForestRegressor’ class of the ‘scikit-learn’ package (Pedregosa et al., 2011).

#### 3.2.3. Neural network architecture

We chose LSTM units (Supplementary data S1) to construct neural network models, as LSTM is a widely used architecture for modelling temporal dependencies in sequential data. A unified neural network structure used for both single-genotype and multiple-genotype prediction is illustrated in Figure 2. All models shared the same temporal backbone, consisting of stacked LSTM layers that process temperature (**T**) and time index (**J**) inputs in a forward pass.

**Figure 2:**
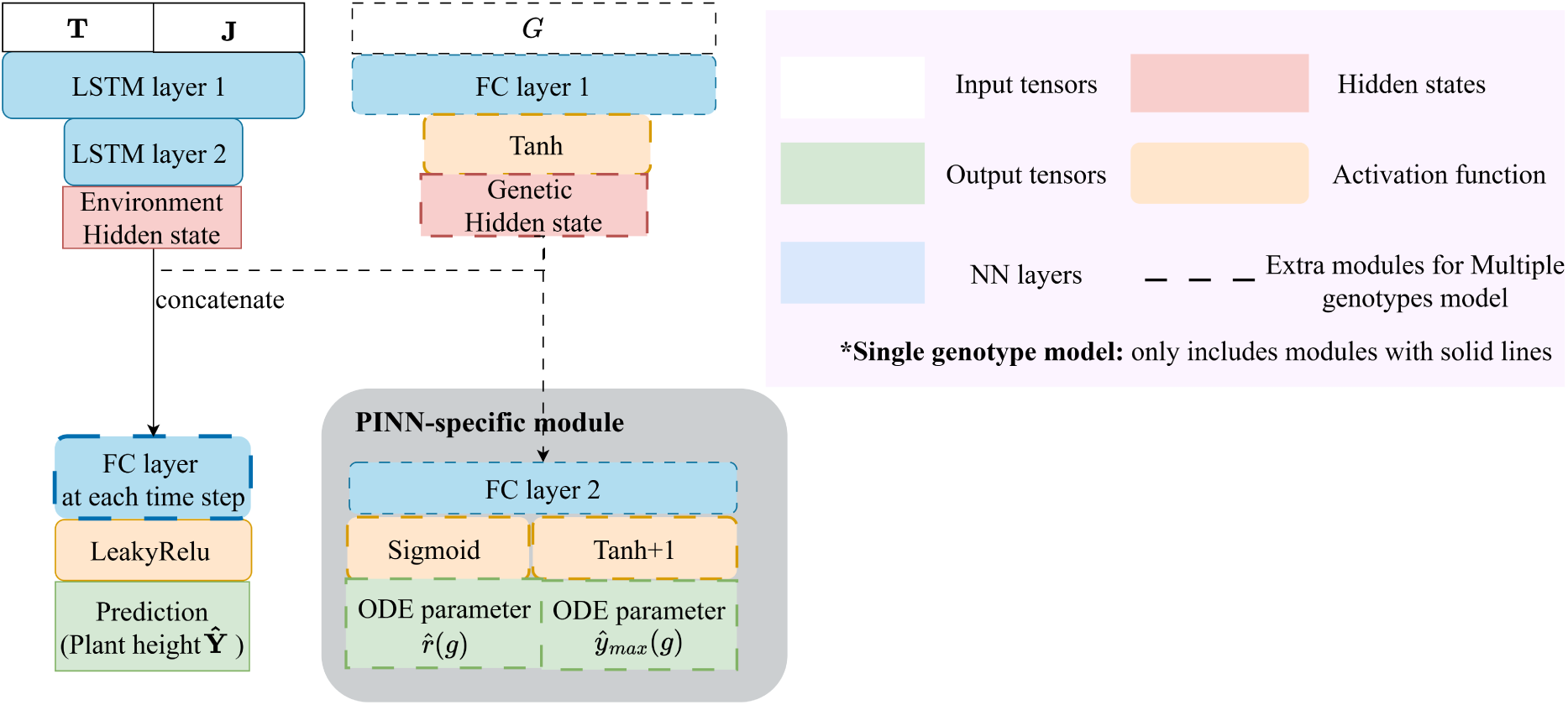
Unified neural network architecture for plant height prediction. The temporal backbone (solid line part) consists of stacked LSTM layers that process temperature (**T**) and time index (**J**) inputs, followed by fully connected (FC) layers and a Leaky ReLU activation to produce plant height predictions *ŷ*. For multiple-genotype models, genotype information **G** was mapped to a latent genetic representation through a genetic embedding module and concatenated with the LSTM hidden state. The grey dashed block denotes the PINN-specific module used only in the *Logi-PINN*, which links genetic information to genotype-specific ODE parameters. In the single-genotype setting, only components shown with solid lines were used.

The output of the LSTM layers was mapped to plant height predictions through fully connected layers followed by a Leaky ReLU activation function. The Leaky ReLU activation constrains predictions to be near-zero or positive values, which is biologically consistent with plant height measurements, and helps mitigate the dead-neuron problem associated with standard ReLU activations (Maas et al., 2013; Xu et al., 2020). In the single-genotype *Logi-PINN*, two additional genotype-specific trainable ODE parameters (*r̂*(*g*) and *ŷ_max_*(*g*)) were used solely to define the derivative-based physics loss during training and were not part of the neural network forward pass that generates plant height predictions. In the single-genotype setting, only components indicated by solid lines in Figure 2 were used.

For multiple-genotype prediction, a genetic embedding module was included, which uses a fully connected layer to map genotype information to a latent genetic representation that was concatenated with the LSTM hidden state before the final fully connected layers. To limit model complexity, genetic effects were assumed to be constant over the growing season; consequently, the parameters of the final fully connected layers that combine genetic and environmental information were shared across all time steps. The *Logi-PINN* extended this shared architecture by introducing a PINN-specific module (grey dashed block in Figure 2) that linked the genetic embedding to genotype-specific ODE parameters used in the physics-based loss formulation. This module was not present in the *LSTM-NN*. Activation functions in this module were selected to enforce biologically meaningful parameter ranges: a sigmoid function constrained the growth rate *r* to values between 0 and 1, a tanh function allowed both positive and negative genetic effects, and a shifted tanh activation (tanh +1) constrained *y*_max_ to the range [0, 2].

#### 3.2.4. Loss components description

We constructed our *Logi-PINN* by integrating a Logistic ODE (Van Voorn et al., 2023) with LSTM (Hochreiter and Schmidhuber, 1997). The LSTM unit was used for the feed-forward pass, following a similar architecture to that described in Section 3.2.3 (Figure 2). The key difference between the *LSTM-NN* and the *Logi-PINN* lies in the loss function, where additional physics-based loss components were introduced. The loss components used for training the *LSTM-NN* and *Logi-PINN* models, and their inclusion under single- and multiple-genotype settings, were summarised in Table 2.

**Table 2.** Loss components used for training LSTM-NN and Logi-PINN models under single- and multiple-genotype settings.

| Loss type | Loss component | LSTM-NN | Logi-PINN<br>(Single-genotype) | Logi-PINN<br>(Multiple-genotype) |
| --- | --- | --- | --- | --- |
| Data-driven | Data loss $L_d$ | ✓ | ✓ | ✓ |
| Regularisation | Weight regularisation $L_2$ | ✓ | ✓ | ✓ |
| Physics-based | Derivative loss: $L_m$ | – | ✓ | ✓ |
| | $r$ penalty loss: $L_r$ | – | ✓ | ✓ |
| | $y_{max}$ penalty loss: $L_y$ | – | – | ✓ |

Note: The LSTM-NN uses the same set of loss components for both single- and multiple-genotype model settings.

We defined the individual loss components as follows. Data loss (*L_d_*) is defined as:

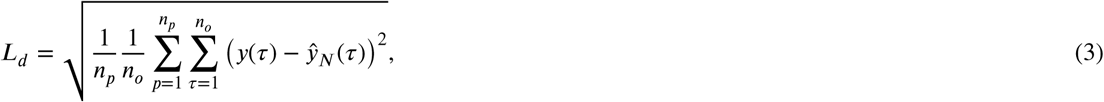

where *ŷ_N_* is the neural network–predicted plant height and *n*_*0*_ < *n_t_* is the number of time points for which plant height measurements were available for each sample. The time indices were defined as *τ* ⊂ {1, 2, …, *n*_*0*_}, where each *τ* corresponds to a day with observed plant height. The time indices were defined as *τ* ⊂ 1, 2, …, *n*_*0*_, where each *τ* corresponds to a day with an observed plant height. Data loss is the key loss for *LSTM-NN* training and focuses on minimising the differences between predicted and observed target variables (plant height). However, this loss function does not impose constraints on intermediate dates (i.e., time points without target trait observations). At those time points, the physics loss serves as the main constraint, since it is calculated at every time step.

The weight regularisation term (a penalty on the squared magnitude of the model’s weights), also known as weight decay, was another loss component used in both the *LSTM-NN* and *Logi-PINN* models. It is defined as:

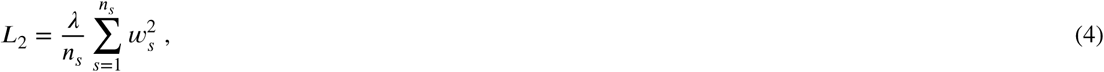

where *λ* is a hyperparameter (a fixed parameter during model training that controls the regularisation strength). The *n*_*s*_ is the total number of weights, *s* indexes the weights and *w*_*s*_ represents the individual weights from the neural network layers. This regularisation term is a commonly used approach to mitigate overfitting by penalising large weight values and reducing less important weights to a small number (Ying, 2019).

The *Logi-PINN* structure introduces physics losses including derivative loss, *r* penalty loss and *y_max_* loss, which can be seen as extra regularisation terms, i.e. an additional penalty based on the difference between the model prediction and a physical law, which is *Logi-ODE*. The *Logi-ODE* parameters (genotype-specific *r*(*g*) and *y_max_*(*g*)) were treated as trainable and are updated either jointly with the neural network weights (single-genotype model) and biases or were predicted based on genetic input features (as illustrated in the grey section of Figure 2). Specifically, the total physics loss is a sum of different loss components: *L*_*m*_, *L_r_*, and *L_y_* loss. The derivative loss (*L*_*m*_) for each sample is defined as:

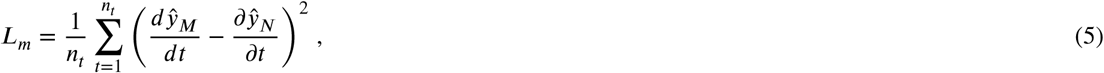

which represents the difference between 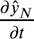, the partial derivative of neural network predicted plant height with respect to time (calculated with auto-gradient from the neural network based on the PyTorch library (Paszke et al., 2019)), and 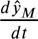 calculated from the mathematical equation (*Logi-ODE*, equation 1). Unlike data loss, the derivative loss was computed at every time step (with a daily resolution). While this could in principle be performed at any temporal resolution, we restricted the model to daily time steps due to limited availability of higher-resolution plant height data and our focus on predictions across the growing season. Importantly, this formulation does not require an explicit initial condition, as we did not integrate the *Logi-ODE* directly; instead, we used the predicted values from the neural network. Hence, 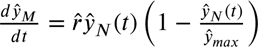.

To ensure that the growth rate estimated by *Logi-PINN* for each genotype (*r̂*(*g*)) remains positive and biologically meaningful (as the overall growth rate parameter *r*(*g*) was assumed to be constant and positive during the growing season), we introduced a penalisation term for negative values of *r̂*(*g*), defined as:

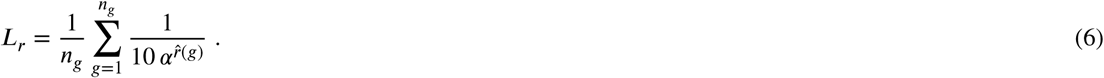

Here, *α* is a fixed scaling parameter controlling the steepness of the penalty. In this study, *α* was set to 10^4^ as a pragmatic choice that ensures a rapidly increasing penalty for negative *r̂*(*g*) values while maintaining numerical stability. This penalty term was chosen because equation 6 is continuously differentiable and strongly discourages biologically implausible negative growth rates. As we estimated *r̂*(*g*) for each genotype *g*, the *L_r_* loss was calculated by averaging across all genotypes in multiple-genotype models. Furthermore, for the multiple-genotype model, the *y*_max_ loss (*L_y_*) helps to reduce the residual between the model-predicted maximum plant height and the genotype-specific ODE parameter *y*_max_ for derivative loss calculation, which was defined as:

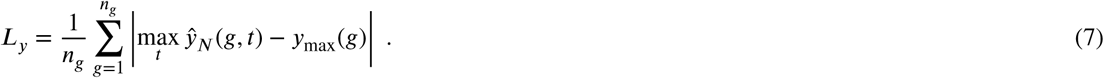

Here, *ŷ*_*N*_ (*g*, *t*) denotes the neural network–predicted plant height at time *t* for genotype *g*, and *y*_max_(*g*) represents the genotype-specific *y*_max_ parameter obtained from the genetic embedding. This loss was applied only in the multiple-genotype model to encourage genotype-specific maximum plant heights and better support genotype-level dynamics in the derivative-based physical constraint.

### 3.3. Model comparison

Models were evaluated under two prediction settings described previously in Section 3.2 and summarised in Table 3. To enable a fair comparison between ODE-based and ML-based models, predictions from the ODE models were generated as follows. Averaged *Logi-ODE* and *Temp-ODE* parameters estimated from the training year were used to generate growth curves, which were then compared with the validation and test year data to calculate prediction performance. For ODE-based and tree-based methods, no hyperparameter tuning protocol was applied, as the results were stable across different optimisation algorithms and model structures (e.g., tree depth and number of trees).

**Table 3.** Prediction settings used for model evaluation.

| Model class | Model | Single-genotype | Multiple-genotype |
| --- | --- | --- | --- |
| ODE-based model | Logi-ODE | ✓ |  |
|  | Temp-ODE | ✓ |  |
| tree-based model | RF | ✓ |  |
| NN-based model | LSTM-NN | ✓ | ✓ |
|  | Logi-PINN | ✓ | ✓ |

For the NN-based models, a common training and evaluation protocol was adopted as described below. We first conducted a coarse exploration of hyperparameters (model depth and width, weight of physics loss) using the training set to identify configurations that allowed stable model convergence. Based on this preliminary exploration, most hyperparameters were fixed to reduce computational cost and to isolate the effects of key design choices. The network architecture was intentionally kept small, with fewer than 300 trainable parameters, to match the limited size of the training dataset and to reduce the risk of overfitting. Hyperparameter tuning was then performed using a small grid search over four discrete hyperparameter combinations, focusing on two parameters based on empirical choice: the hidden size (3 or 5) of the first LSTM layer output and the weight of the physics-based loss (2 or 9). The optimal hyperparameter configuration was selected following the procedure described below and used for model evaluation and comparison.

Model weights were initialised before training using orthogonal matrices, which have been shown to improve performance and stability in recurrent neural networks (Saxe et al., 2013). Masked RMSE (Equation 3), computed on the validation set, was used as the criterion for early stopping, hyperparameter tuning, and model comparison. Models were trained for a maximum of 3000 epochs, and checkpoints were saved every ten epochs. Final model selection was performed by identifying the checkpoint with the minimum validation RMSE after 1500 epochs based on the validation set. This stopping criterion was chosen to avoid overfitting while also preventing premature termination due to uncertainty in weight initialisation and large fluctuations during the early stages of training (Frankle et al., 2020). Additionally, each hyperparameter combination was trained five times with different random seeds as a practical compromise between computational cost and stability assessment. For each combination, the mean and standard deviation of the validation RMSEs were calculated, and the hyperparameter combination with the minimum average validation RMSE was selected for model comparison.

As we focused on results obtained from a selected data split to improve training data diversity (Section 3.1.4), we additionally reported results from another five splits (six in total) using different combinations of training years in supplementary data S4. For each split, the average RMSE on the test set was computed for both the *LSTM-NN* and *Logi-PINN* models.

## 4. Results

### 4.1. Single-genotype model plant height curve prediction

Table 4 shows the average RMSE scores for the training (two years), validation (one year), and testing (one year) sets of the five models covering the 19 genotypes. The standard deviation (sd) values for each model were determined from five different runs per genotype for each separate model, with each run using a different random parameter initialisation (by using different random seeds for initialisation).

**Table 4.** Five single-genotype model RMSE comparision.

| Model | Training RMSE (±sd) | Validation RMSE (±sd) | Test RMSE (±sd) |
| --- | --- | --- | --- |
| *Logi-ODE* | $0.045 \pm 0.000^a$ | $0.070 \pm 0.000$ | $0.111 \pm 0.000$ |
| *Temp-ODE* | $0.108 \pm 0.001$ | $0.063 \pm 0.002$ | $0.103 \pm 0.004$ |
| RF | $0.020 \pm 0.002$ | $0.133 \pm 0.000^a$ | $0.188 \pm 0.000^a$ |
| *LSTM-NN* | $0.032 \pm 0.008$ | $0.056 \pm 0.011$ | $0.072 \pm 0.022$ |
| *Logi-PINN* | $0.029 \pm 0.004$ | $0.060 \pm 0.007$ | **$0.057 \pm 0.008$** |

This table shows the RMSE averaged results from single-genotype models (average across 19 genotypes) with selected two distinct training years, 2018 and 2019. Our *Logi-PINN* was the best performing model based on test RMSE.
<sup>a</sup> sd is 0.000 after rounding to three decimal places.

In most cases, the *Logi-ODE* recreated the shape of the growth curve (figures can be found in Supplementary Figure S5.1) but did not provide accurate predictions: the test RMSE of the *Logi-ODE* was approximately twice that of the best-scoring model, i.e. the *Logi-PINN*. The test RMSE of the *Temp-ODE* was only slightly better than that of the *Logi-ODE*. The training involved two years of data with different temperature profiles. However, in most cases, the predicted curves from the *Temp-ODE* failed to distinguish between the two years, as indicated by the high training RMSE. The *RF*yielded the worst validation and test RMSE values. This was likely the result of overfitting, given that the training RMSE of the *RF* was the lowest among all five models and the sd for the test RMSE was very small. The *LSTM-NN* had the second-lowest test RMSE, and its validation RMSE was the lowest among all five models. However, the sd values of the *LSTM-NN* were the largest among all five models. The *Logi-PINN* achieved the best overall performance: its test RMSE was the lowest among all models, while the corresponding standard deviation was relatively small. Moreover, the parameters *ŷ_max_* and *r̂* of the *Logi-PINN* across different runs consistently converged to similar values after training, which further indicates the stability of the model (see Supplementary Figure S6.1).

To visually compare the predicted plant height curves, we plotted predicted plant height (curves) and measured plant height (dots) from our *LSTM-NN* and *Logi-PINN* models in Figure 3. The figure shows two example genotype prediction results in the test set, in which the shade (area around the solid line) indicates the interval (mean ± sd) calculated from different model parameter initialisation. The *LSTM-NN* and *Logi-PINN* accurately predicted plant height growth curves over time, which is consistent with the established capacity of LSTM architectures to capture local temporal dynamics. Specifically, the predicted curves followed the same overall shape as measured plant height, for instance, the observed bump at around 100 days in dots is also captured by the predicted curves (Figure 3). However, predicting the plant height in the middle of the growing season was more challenging (the slope is large), particularly when the plants were growing rapidly and were likely more sensitive to changing environmental factors. The *Logi-PINN* had smaller fluctuations compared to *LSTM-NN* (Figure 3). In particular, after plants reached their maximum height (after day 100), the *LSTM-NN* predictions continued to vary over time, whereas plant height was expected to remain stable.

**Figure 3:**
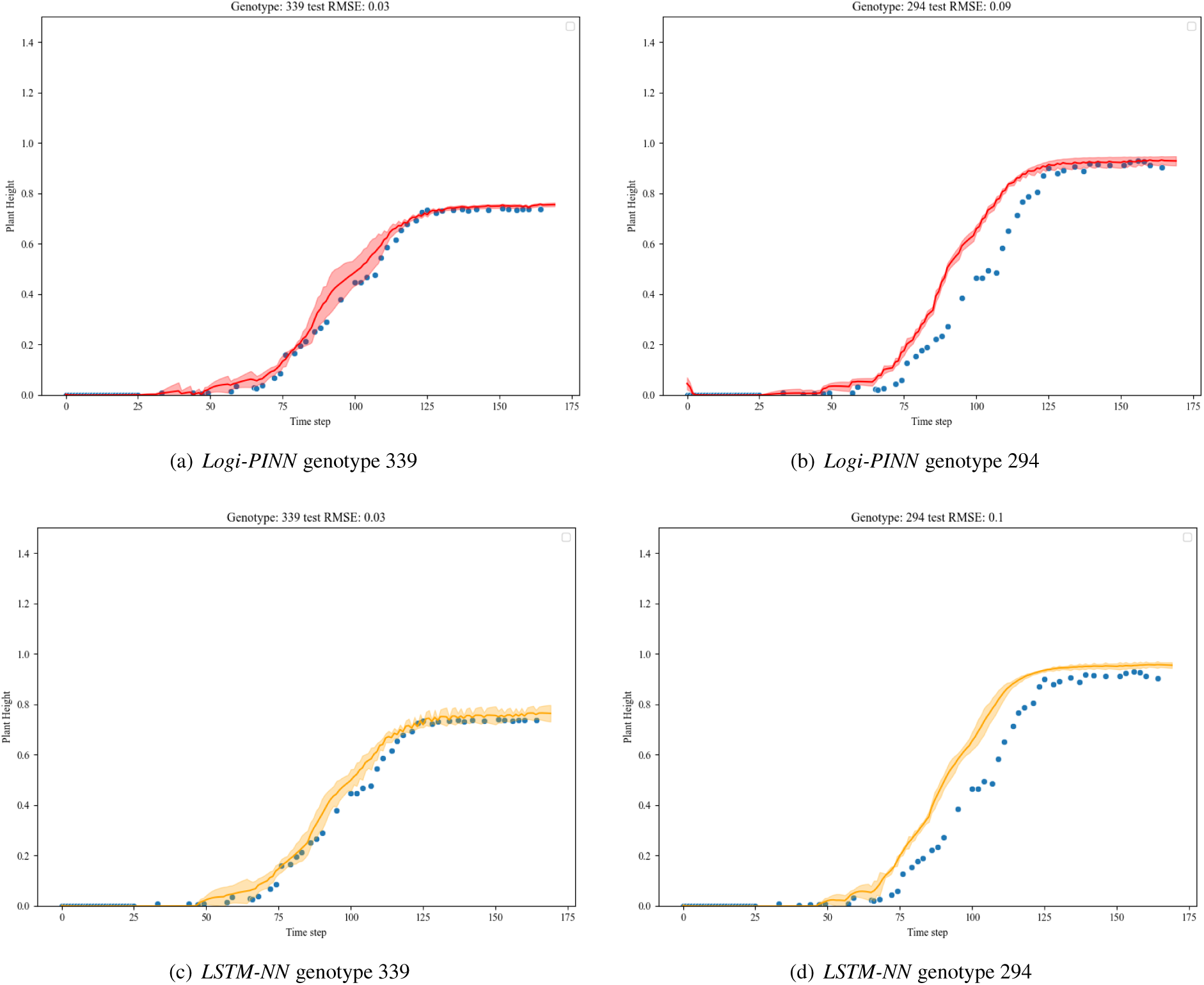
Predicted and observed plant heights for two models and two genotypes in the test set. The *Logi-PINN* (top) and *LSTM-NN* (bottom) results for genotype 339 are shown as an example of a well-fitted genotype, while genotype 294 is shown as an example of a less well-fitted case. The blue dots represent the measured data, and the orange and red curves represent the averaged predicted plant height. The shaded area represents the interval (mean ± sd) at each time step, calculated from model-predicted plant height using five different parameter initialisations. The *LSTM-NN* results tend to have larger sd values and less smooth predictions.

To further investigate the performance of our models, we also compared the models’ test RMSE across genotypes (Figure 4). We can see there is a variation in test RMSE between genotypes. While the *Logi-PINN* does not always have a lower test RMSE compared to the *LSTM-NN*, the variance of the test RMSE was consistently smaller for the *Logi-PINN* than for the *LSTM-NN* (orange dots spread wider than red dots). The other three models had a smaller sd, while they had a higher test RMSE for most genotypes.

**Figure 4:**
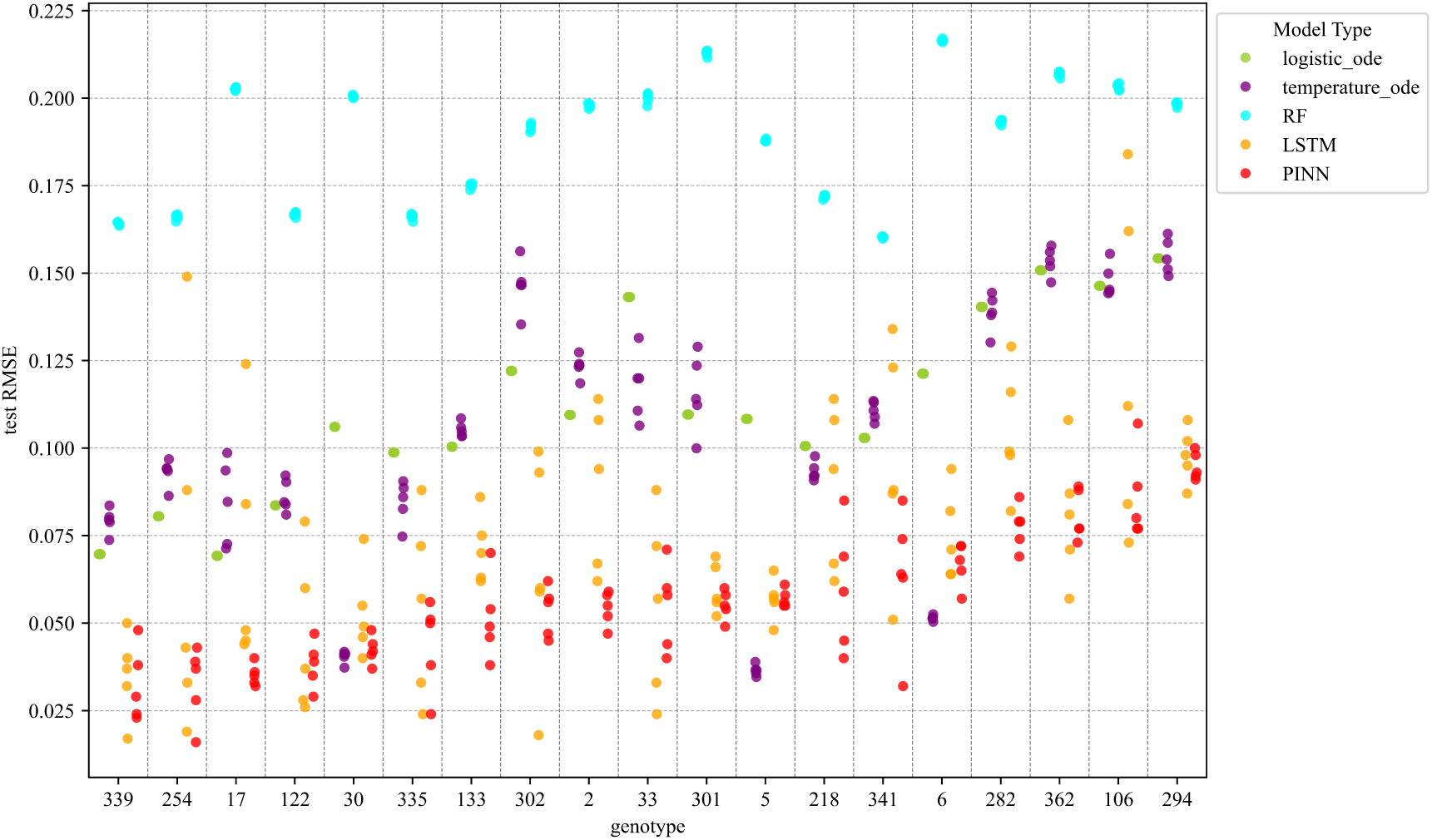
RMSEs for individual genotypes for different single genotype models. Each genotype was separated by the dashed lines and in order of PINN RMSEs. A dot is a test RMSE for a single initialisation of a single model applied to a single genotype.

### 4.2. Multiple-genotype model plant height curve prediction

Although different genotypes had different prediction accuracies, the overall growth curve shape was still similar, as we can see in Figure 3. Table 5 shows the result of multiple-genotype models for unseen year prediction (Year split), where we wanted to test if we can improve the prediction performance by borrowing information from other genotypes.

**Table 5.** RMSE results for multiple-genotype models based on year splits. The text in bold was the minimum test RMSE score for each encoding method.

| Split Methods | Models | one-hot encoding |  |  | Kinship Matrix Encoding |  |  |
| --- | --- | --- | --- | --- | --- | --- | --- |
|  |  | Training RMSE | Validation RMSE | Test RMSE | Training RMSE | Validation RMSE | Test RMSE |
| Year Split | *LSTM-NN* | $0.035 \pm 0.001$ | $0.076 \pm 0.010$ | **$0.058 \pm 0.012$** | $0.037 \pm 0.004$ | $0.073 \pm 0.007$ | **$0.059 \pm 0.007$** |
| | *Logi-PINN* | $0.034 \pm 0.002$ | $0.070 \pm 0.009$ | $0.061 \pm 0.007$ | $0.037 \pm 0.006$ | $0.067 \pm 0.006$ | $0.065 \pm 0.017$ |

The result shows that the multiple-genotype *Logi-PINN* had a higher test RMSE than the single-genotype model, while the multiple-genotype *LSTM-NN* had a lower RMSE than the single-genotype model setup. The decrease in test RMSE for the *LSTM-NN* model could be attributed to the increased amount of training data. The model was trained using a dataset that was 19 times larger than that of a single-genotype model, as data from all genotypes were used to train a single model. This allowed the LSTM layers to extract shared information across genotypes and predict plant height under different temperature conditions for the given genotypes. While for the *Logi-PINN* model, the only difference in the model’s structure was that we linked genotypes with ODE parameters used for *Logi-PINN* training. This encoding structure implied a relationship between ODE parameters and genetic information, which was not clear from our current dataset; further information can be found in our discussion section.

We also plotted the predicted curves of two representative genotypes (good and bad prediction performance example). The sd for *Logi-PINN* and *LSTM-NN* was relatively small during the growing season. There was a clear improvement in the stability of *LSTM-NN* with a multiple genotype setup (Figure 3 c, d and Figure 5 c, d).

**Figure 5:**
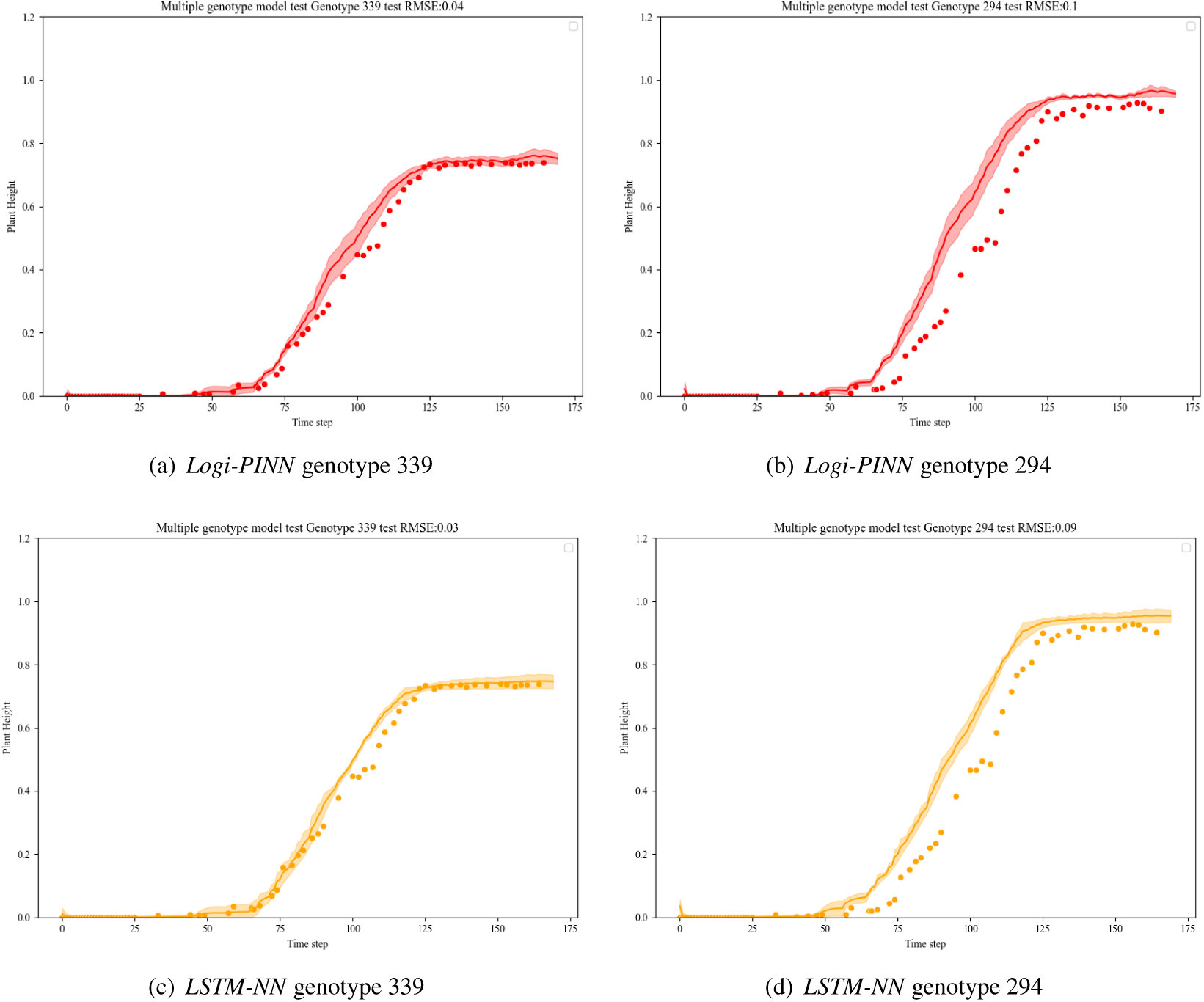
Predicted and observed plant heights for the multiple-genotype model (kinship encoding as an example). *LSTM-NN* model became more stable and got a smoother result than the single-genotype setup. There was no gain for the *logi-PINN* model with multiple genotype setups. The uncertainty of prediction came from different model initialisations, which mainly affected prediction in the middle of the growing season and at the very end.

## 5. Discussion

We discuss the observed differences in model performance with reference to the advantages and limitations summarised in Table 6.

**Table 6.** Advantages and disadvantages of four selected model structures.

| Model | Advantages | Disadvantages |
| --- | --- | --- |
| ODEs (Process-based models) | 1. Consider time dependency (Teschl, 2024). Include expert knowledge of system development, thus making it easy to interoperate (Polynikis et al., 2009) | 1. Difficult to fit when feature space and parameter space increase (Wu et al., 2019; Willms, 2007). |
| RF (Classical ML model) | 1. Robustness to hyperparameter choice (Probst et al., 2018).<br>2. Suitable for high-dimensional data (Ghosh and Cabrera, 2021; Schwarz et al., 2010). | 1. Does not capture time dependency or requires specific feature design (Regier et al., 2023).<br>2. The prediction is bound to the range of training data and is not sufficient for prediction in a new range (Breiman, 2001). |
| LSTM-NN (Neural network model) | 1. LSTM is specifically designed to address gradient vanishing in long time series, which allows representation of various time-lag effects (Dupont et al., 2024; Hochreiter and Schmidhuber, 1997).<br>2. A flexible structure suitable for end-to-end training (Sukhbaatar et al., 2015; Wang et al., 2019) allows the model to train and perform feature selection based on its end task (plant height curve prediction). | 1. More parameters are included compared to process-based models, which makes the model less interpretable (Liang et al., 2021).<br>2. No explicit domain knowledge is encoded. |
| Logi-PINN (PINN model) | 1. Captures time dependency and time-lag effects (Nathasarma and Roy, 2023; Hochreiter and Schmidhuber, 1997).<br>2. Physics loss can constrain model predictions where label data are not available, constraining the solution space (Hu et al., 2024; Karniadakis et al., 2021).<br>3. Suitable when fewer data are available and when given physical laws are followed (Blechschmidt and Ernst, 2021). | 1. Difficulty in balancing different parts of the loss function (Cuomo et al., 2022; Nandi et al., 2022).<br>2. Prediction accuracy and interoperability also depend on the reliability of the ODE (physics) component. |

### 5.1. LSTM-based model outperforms other compared methods in prediction accuracy

Our results suggest that our LSTM-based models, with their ability to capture temporal dependencies, can accurately predict plant height during the growing season using only air temperature. The *Logi-PINN* achieved the lowest test RMSE in the single-genotype experiments, outperforming all other models. The *LSTM-NN* improves when more training data are available in the multiple-genotype setup. The best-performing *Logi-PINN* and the best-performing *LSTM-NN* showed comparable accuracy in the multiple-genotype setting.

The RNN structure was chosen because it is biologically meaningful in terms of processing time and temperature inputs sequentially. The LSTM unit takes temperature inputs from both the current and previous time steps, which is comparable to the growing degree day concept used in crop modelling (McMaster and Wilhelm, 1997). It can accumulate temperatures from previous days through its recurrent hidden states, summarising past temperature information to influence growth predictions. Using a recurrent structure to process raw temperature input instead of calculating growing degree days is more flexible, as it uses different weights that adjust how much information is retained or forgotten from previous time steps to make the current prediction. The weights of different gates are adjusted based on the input data (an example of gate weights from the first LSTM layer of the *Logi-PINN* model is shown in Figure S7.1). In contrast, *RF* is a commonly used data-driven model in agriculture but performs poorly due to its known limitations in time-series tasks compared to dynamic models. Thus, *RF* models used for time-series prediction and classification often rely on either manually designed features to capture temporal trends (Karasu and Altan, 2019) or the use of previous values of the predicted variable to introduce temporal dependencies (Tyralis and Papacharalampous, 2017).

Process-based models, such as the ODE approach used here, are built on interpretable mathematical formulations with a limited number of parameters. These models are an oversimplified summary of real-life systems. The simplified plant growth assumptions and limited number of parameters in the ODE structure restrict the flexibility and capacity of ODE models. In our case, the logistic structure restricts the growth curve shape and assumes a constant basic growth rate throughout the season. Even with *Temp-ODE*, we assume that air temperature has a direct effect on growth rate. The current ODE structure is less effective in tracking finer details of plant height fluctuations arising from more complex effects, such as lagged environmental effects on growth rate. Adding additional ODEs that link other environmental responses to plant height could increase flexibility, but this requires accurate domain knowledge about the system and increases optimisation complexity. Furthermore, more complex process-based models designed to describe detailed mechanisms of plant growth are difficult to calibrate and increase uncertainty in predictions (Droutsas et al., 2022; Dokoohaki et al., 2021). The choice of equations and calibration of parameters also strongly influence model performance and often rely on user expertise (Confalonieri et al., 2016).

PINN structures provide a way to integrate known complex biological processes while retaining the expressive power of deep learning, making them especially suitable for modelling dynamic plant traits such as growth over time. Even with a simple *Logi-ODE* describing the global trend of plant height over the growing season, prediction accuracy can be improved with limited data. The additional loss terms allow the model to incorporate multiple criteria simultaneously during training, which may improve the calibration process compared to single-objective optimisation, as discussed in process-based modelling (Wagener et al., 2003). Furthermore, adding physics loss reduces uncertainty across different model initialisations and yields more stable predictions, as reflected by decreased sd compared to *LSTM-NNs*. The larger sd observed in the single-genotype *LSTM-NN* results also suggests a higher tendency to overfit noisy patterns in the training data, thereby reducing generalisation ability.

### 5.2. Potential use and future study of the temporal plant growth prediction model

While our current temporal prediction study focused on plant height, the same model structure could be adapted to forecast other dynamic plant traits, such as leaf area, canopy cover, or biomass accumulation. The HTP platform produces phenotyping data from different plant organs, revealing dynamic plant development (Tardieu et al., 2017), and these data can be used to train multi-trait temporal prediction models. Building on this, a well-performing PINN model that accurately forecasts multiple temporal traits can serve as a biologically meaningful basis for yield prediction. Specifically, it can be extended to yield prediction by decomposing yield into underlying components or intermediate traits (Tsutsumi-Morita et al., 2021). Instead of treating yield as a static prediction task, dynamic prediction of multiple traits can be combined to inform yield prediction. In this way, predictions reflect the logical development of intermediate traits and can be constrained by intermediate trait dynamics.

The potential use of a temporal prediction model lies in converting temporal phenotyping data from the HTP platform and static genetic data into genetic gain (Araus et al., 2018; Roth et al., 2023), supporting selection decisions that accelerate plant breeding. Combining data-driven and process-based models to embed marker data and link them to temporal traits provides a potential way to extend prediction models to new cultivars. Machine learning methods are becoming increasingly important for analysing genetic data (Libbrecht and Noble, 2015), and more explainable ML models are essential for understanding genetic effects (Novakovsky et al., 2023). PINN and other hybrid ML methods can not only reduce data requirements but also improve interpretability, as domain knowledge and appropriate rules are encoded in the model structure or guide the model learning process.

### 5.3. Current limitations of PINN for plant growth systems

To support the further development of PINNs for dynamic plant growth systems, it is important to carefully consider challenges associated with their application to real-life biological systems. Previous PINN studies have mainly focused on controlled engineering systems or well-established physical systems, where integrated equations provide near-perfect descriptions of underlying processes (Linka et al., 2022; Wong et al., 2025). However, real-life systems such as plant growth introduce additional challenges for PINNs: they are often more complex than the domain knowledge summarised by experts in process-based models (Camargo and Kemanian, 2016), and non-linear environmental noise can complicate PINN optimisation (Pilar and Wahlström, 2024).

Addressing these challenges requires careful consideration when designing model structures for real-life systems. Our results suggest that inaccurate prior information can hinder model performance more severely than the absence of such information. For example, in the multiple-genotype setting, inaccurate information may arise from assumptions of direct relationships between ODE parameters and genotype similarity used in constrained training procedures. As genotype distance is incorporated as an additional constraint during training, inaccuracies in genetic similarity representations are likely to have a larger impact on PINN-based models than on purely data-driven approaches. In this study, SNP information was summarised into a single similarity measure, which provides a simple representation that facilitates unseen-year prediction but may lack sufficient information to support extrapolation to unseen genotypes (Supplementary Section S8). Consistent with this observation, no clear relationship was found between ODE parameters and genotype similarity (Figure S8.1). This helps explain why the *Logi-PINN* exhibits lower prediction accuracy compared to the single-genotype setup.

In contrast, less informative but soft constraints are likely to cause less harm to model performance (Márquez-Neila et al., 2017). Although the *Logi-ODE* component in the current *Logi-PINN* structure does not explicitly capture environment-related growth variations, it still leads to more stable and accurate predictions in the single-genotype setting. As a soft constraint, it does not introduce strong structural bias, allowing environmental effects to be learned directly from input temperature data. This suggests that partially correct physical rules, which may not fully describe system dynamics, can be incorporated as soft constraints to maintain model flexibility while avoiding restrictive assumptions. This consideration is particularly important for real-life biological systems such as plants, where physical rules are often derived under assumptions that may not always hold.

Another important consideration is how PINN training balances multiple components in the loss function, typically combining data loss and physics loss, resulting in a multiobjective optimisation problem (Gunantara, 2018). In this study, the weight of the physics loss was treated as a hyperparameter and searched over a discrete space, introducing two potential challenges. First, different loss components may conflict with each other, particularly when physics-based models do not fully capture the complexity of real-world systems, leading to discrepancies between process-based assumptions and observed data (Xu et al., 2023). Second, even when loss components do not drive optimisation in opposing directions, their relative scales can vary during training, causing the model to focus disproportionately on one component. Allowing dynamically adjusted loss weights during training may help alleviate this issue (Gao et al., 2025). Previous studies suggest that *L*_2_ regularisation on neural network parameters or alternative regularisation strategies for physics loss can improve PINN stability and prevent dominance of either data loss or physics loss during training (Pu and Chen, 2023; Wang et al., 2022).

## 6. Conclusion

Environmental factors dynamically influence plant growth throughout the growing season rather than directly determining the end-season phenotype. Capturing these temporal dynamics is essential for accurately predicting plant growth under new environmental conditions. We propose a *Logi-PINN* structure that incorporates dynamic effects by combining a time-dependent neural network with an ODE structure to predict plant height for different genotypes under new environmental conditions. This work demonstrates the applicability of PINN frameworks for temporal plant growth trait prediction. The results show improved average prediction accuracy and robustness when physics loss is incorporated in single-genotype models. This PINN structure is adaptable to multiple dynamic phenotypes when appropriate data are available. The proposed temporal prediction model can bridge the gap between dynamic phenotype prediction and genotype information and help shorten breeding cycles.

## Supporting information

All supplementary files

## Abbreviations

**Table 1.** List of abbreviations used in this manuscript.

| Abbreviation | Definition |
| --- | --- |
| DE | Differential Equation |
| FIP | Field Phenotyping Platform |
| FC | Fully Connected |
| HTP | High-Throughput Phenotyping |
| LSTM | Long Short-Term Memory |
| ML | Machine Learning |
| NN | Neural Network |
| ODE | Ordinary Differential Equation |
| PINN | Physics-Informed Neural Network |
| RF | Random Forest |
| RMSE | Root Mean Squared Error |
| RNN | Recurrent Neural Network |
| SNP | Single-Nucleotide Polymorphism |
| Temp-ODE | Temperature-informed Ordinary Differential Equation |
| Logi-ODE | Logistic Ordinary Differential Equation |
| Logi-PINN | Logistic Ordinary Differential Equation–Informed Physics-Informed Neural Network |

## 7. Supplementary data

All supplementary data are included in one PDF file and organised into separate sections based on topic. The following supplementary data are available at JXB online.

Section S1: Includes Figure S1.1, which provides a brief description of the LSTM unit used to construct our models.

Section S2: Includes Notation Table S2.

Section S3: Includes Figure S3.1 and Figure S3.2, which illustrate an example of the HTP dataset and the daily temperature data used in this study.

Section S4: Includes Figure S4.1 and Table S4, showing single-genotype cross-validation results for the *LSTM-NN* and *Logi-PINN* models.

Section S5: Includes Figure S5.1, which shows *Logi-ODE* fits for each replicate (plot) separately.

Section S6: Includes Figure S6.1, which shows the convergence stability of *Logi-PINN* parameters.

Section S7: Includes Figure S7.1, which shows weight parameters from the trained PINN model for different genotypes.

Section S8: Includes Figure S8.1, which shows the relationship between genotype similarity and *Logi-ODE* parameters.

## 8. Acknowledgements

Thanks to helpful discussions with Lukas Roth and Andreas Hund on the dataset.

## 9. Author contributions

**Yingjie Shao:** Conceptualisation; Methodology; Software; Writing – original draft **Fred van Eeuwijk:** Supervision; Conceptualization; Writing – review & editing; Funding acquisition **Carel F.W. Peeters:** Supervision; Writing – review & editing; Funding acquisition **Olivia Zumsteg:**Data Curation; Writing – review & editing **Ioannis N. Athanasiadis:** Supervision; Methodology; Writing – review & editing; Funding acquisition **George van Voorn:** Supervision; Methodology; Writing – original draft; Funding acquisition

## 12. Data Availability

The data used in the study is downloaded from Roth et al. (2025). The code and processed data for model training can be found in the Github repository.

