## Supplementary material for "Physics-Informed Neural Network Methods for Predicting Plant Height Development": All supplementary files

### Supplementary Data

#### S1. LSTM unit

As a commonly used neural network for sequential data processing, the LSTM unit includes two states to keep long and short-term memory from time series (Figure S1.1 modified based on (Zhang et al., 2021). The cell state is designed to capture long-term memory in the time series, and the hidden state is strongly related to short-term memory. When the $X_t$ is fed into an LSTM unit, the input from current time step ($X_t$) and Hidden state from last time ($H_{t-1}$) are used to calculate the forget gate $F_t$, input gate $I_t$, output gate $O_t$ and candidate cell state $\tilde{C}_t$ with Equation S1.1 given as

$$\begin{aligned} F_t &= \sigma(X_t W_{xf} + H_{t-1} W_{hf} + b_f) \\ I_t &= \sigma(X_t W_{xi} + H_{t-1} W_{hi} + b_i) \\ O_t &= \sigma(X_t W_{xo} + H_{t-1} W_{ho} + b_o) \\ \tilde{C}_t &= \tanh(X_t W_{xc} + H_{t-1} W_{hc} + b_c). \end{aligned} \tag{S1.1}$$

The calculation assigns separate weights $W_{xf}$,$W_{xi}$ and $W_{xo}$ for current time step input and $W_{hf}$ $W_{hi}$ and $W_{ho}$ for previous hidden state output. The $\sigma$ is a sigmoid activation function used for converting the value between zero and one, which determines whether the corresponding information needs to be forgotten or kept. *tanh* is a hyperbolic tangent activation function that narrows the output values between -1 and 1, with negative values representing negative effects and positive values representing positive effects, respectively. After that, the cell state $C_t$ and hidden state $H_t$ at time t will be updated based on

$$\begin{aligned} C_t &= F_t \odot C_{t-1} + I_t \odot \tilde{C}_t \\ H_t &= O_t \odot \tanh(C_t), \end{aligned} \tag{S1.2}$$

where the $\odot$ notation represents the element-wise multiplication (Hadamard product). Notice that the notations for this section are commonly used in literature, which do not overlap or are used in other parts of this paper.

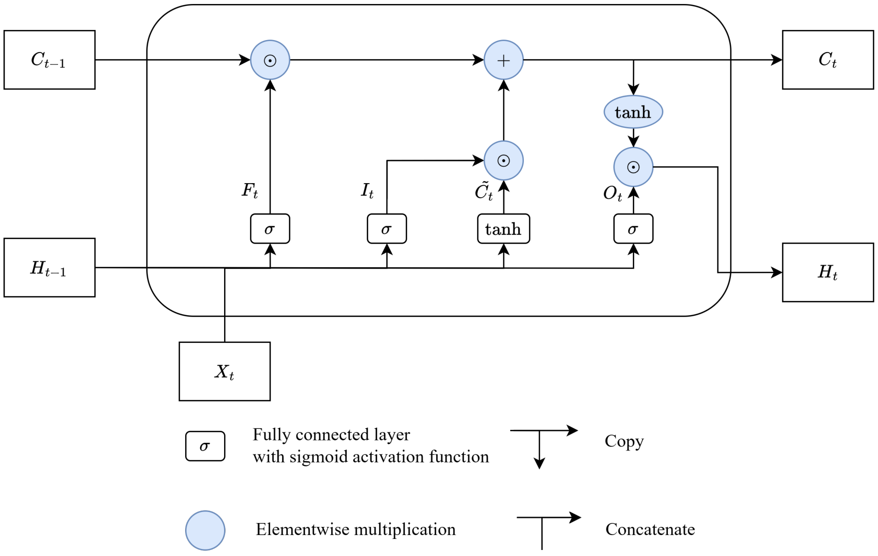


**Figure S1.1:** LSTM unit

#### S2. Notation Table

| Symbol | Description | First use of the symbol |
|---|---|---|
| | **Tensors** | |
| $\mathbf{G}$ | A tensor of Genotype encoding. | section 3.1.3 |
| $\mathbf{J}$ | A tensor of time index. | section 3.2 |
| $\mathbf{T}$ | A tensor of air temperature data (2 meters above ground). | section 3.2 |
| $\mathbf{Y}$ | A tensor of Plant height data. | section 3.2 |
| $\hat{\mathbf{Y}}$ | A predicted plant height tensor, which has the same shape as $\mathbf{Y}$ . | Figure 2 |
| | **Variables & Functions** | |
| $\frac{dy_M(t)}{dt}$ | The derivative of plant height with respect to time $t$ from *Logi-ODE* [a]. | Equation 1 |
| $\frac{d\hat{y}_M}{dt}$ | The derivative calculated based on the mathematical equation (*Logi-ODE*). | Equation 5 |
| $\frac{dy(t)}{dt}$ | The derivative of plant height with respect to time $t$ from *Temp-ODE*. | Equation 2 |
| $\hat{y}_N$ | Neural network predicted plant height [b]. | Equation 3 |
| $\frac{\partial \hat{y}_N}{\partial t}$ | The gradient of predicted plant height with respect to time index calculated from neural network part based on auto-gradient. | Equation 5 |
| $L_2$ | The weight regularisation loss, which is $L_2$ regularisation on neural networks parameters. | Equation 4 |
| $L_d$ | The data loss, an averaged RMSE across the whole time series. | Equation 3 |
| $L_m$ | The derivative loss, which minimises the difference between auto-gradient and ODE derivative. | Equation 5 |
| $L_r$ | The penalization loss, which penalize negative $r$ value during *Logi-PINN* training. | Equation 6 |
| $L_y$ | The $y_{max}$ loss, which reduces the residual between model predicted maximum plant height and genetics embedded $y_{max}(g)$. | Equation 7 |
| $\hat{r}(g)$ | The parameter $r$ for genotype $g$ estimated by *Logi-PINN* models. | Equation 6 |
| $u(t)$ | Temperature response curve that modify growth rate $r$ in *Temp-ODE*. | Equation 2 |
| $y_M(t)$ | The plant height at time step $t$, a state variable of *Logi-ODE*. | Equation 1 |
| $y(t)$ | The plant height at time step $t$, a state variable of *Temp-ODE*. | Equation 2 |
| $\hat{y}_{max}(g)$ | The $y_{max}$ predicted for genotype $g$ from on *Logi-PINN* models. | Equation 7 |
| $\hat{y}_N$ | Neural network predicted plant height. | Equation 3 |
| $\hat{y}_N(g,t)$ | The plant height predicted by NN-based model at time $t$ for genotype $g$. | Equation 7 |
| | **Fixed numbers** | |
| $n_t$ | The total number of days in a growing season in our dataset, in this case $n_t = 170$. | section 3.1.2 |
| $n_o$ | The total number of measured plant height during a growing season, which is around 40 but varies across different year and genotypes | Equation 3 |
| $n_p$ | The number of samples (plots) in the dataset. | section 3.1.2 |
| $n_g$ | The number of genotypes in the dataset. | section 3.1.3 |
| $n_s$ | The total number of weight. | Equation 4 |
| | **Parameters** | |
| $r$ | A parameter from *Logi-ODE* and *Temp-ODE* that describes an overall growth rate across the growing season. | Equation 1 |
| $T_{AL}$ | Arhenius temperature for the rate of decrease at the lower boundary of the temperature tolerance range, which was set as consent (2000 K). | Equation 2 |
| $T_{AH}$ | Arhenius temperature for the rate of decrease at the upper boundary of the temperature tolerance range, which was set as consent (60000 K). | Equation 2 |
| $T_H$ | Upper boundary of the temperature tolerance. | Equation 2 |
| $T_L$ | Lower boundary of the temperature tolerance. | Equation 2 |
| $w_s$ | The individual weights from the neural network layers. | Equation 4 |
| $y_{max}$ | A parameter from *Logi-ODE* and *Temp-ODE* that describes the maximum boundary of plant height. | Equation 1 |
| $\lambda$ | A weight parameter for the regularisation strength. | Equation 4 |
| | **Indices** | |
| $g$ | Index of genotypes $g \in \{1, 2, \ldots, 19\}$ . | Equation 6 |
| $p$ | Index of plots (samples). | Section 3.2 |
| $s$ | Index of weight parameters in neural network layers. | Equation 4 |
| $t$ | Index of time points, and $t = 1, 2, ...n_t$. | Section 3.1.2 |
| $\tau$ | The time indices are defined as $\tau = 1, 2, \ldots, n_t$ (where each $\tau$ corresponds to an observed day for each sample). | Equation 3 |

[a]The subscript $_M$ (e.g., $\frac{dy_M(t)}{dt}$) indicates that the value is obtained from a mathematical model (ODE).
[b]The subscript $_N$ (e.g., $\hat{y}_N$) indicates that values are obtained from an NN-based model (*LSTM-NN* or *Logi-PINN*) or functions related to neural network models.

### S3. Example data

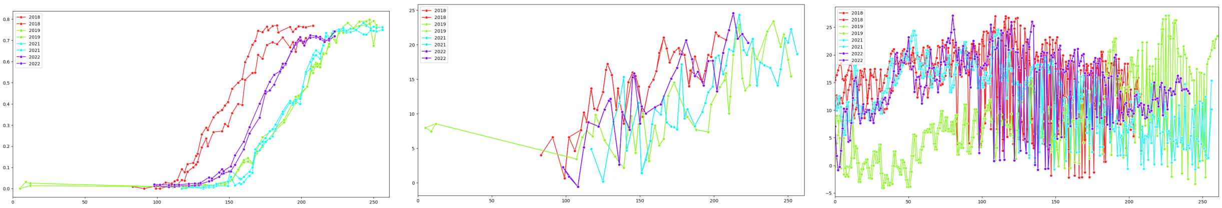

(a) Plant height (b) Temperature (c) Daily temperature

**Figure S3.1:** An example of measured wheat height and corresponding temperature during the growing season. The left figure shows plant height for genotype 335 in four years. The colours correspond to different years, with two replicates within each year. The middle figure is the corresponding daily average air temperature, but at the same time step as measured plant height, and the right figure is for the daily temperature of the whole growing season.

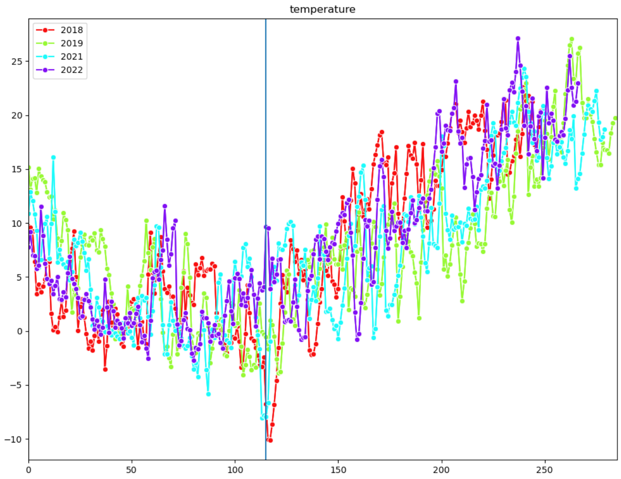


**Figure S3.2:** Daily temperature after alignment based on plan sowing date. The plot shows four-year daily temperature data, which indicated by four colour. The blue vertical line indicates the start date (115 days after sowing) used in our model training.

#### S4. Different data split result for *LSTM-NN* and *Logi-PINN*

**Table S4.1**
Summary of model comparison result for single-genotype model with different data splits.

| Model | Training RMSE $\pm$ sd | Validation RMSE $\pm$ sd | Test RMSE $\pm$ sd |
|---|---|---|---|
| *LSTM-NN* | $0.026 \pm 0.007$ | $0.115 \pm 0.037$ | $0.101 \pm 0.035$ |
| *Logi-PINN* | $0.026 \pm 0.003$ | $0.094 \pm 0.021$ | **0.083** $\pm 0.014$ |

This table shows the average result from single-genotype models with six possible data splits. The standard deviation (sd) was calculated based on random seeds and averaged across different data splits.

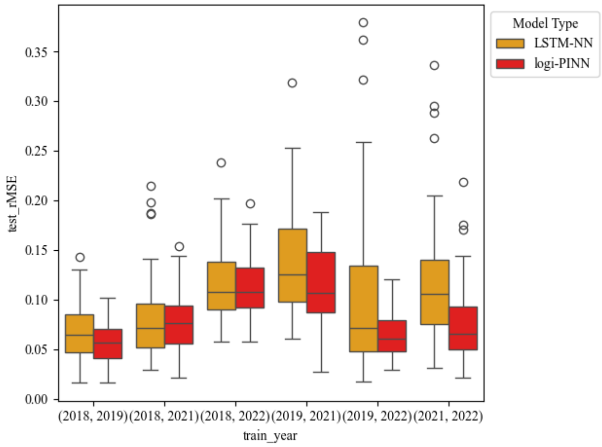


**Figure S4.1:** Different training year *Logi-PINN* and *LSTM-NN* test RMSE result comparison. The box plot shows the test RMSE result for two single-genotype models: *Logi-PINN* and *LSTM-NN*. The models have different prediction accuracy, likely because of the data diversity in the training year. In general, the *Logi-PINN* model has lower test RMSE than the *LSTM-NN* model.

#### S5. *Logi-ODE* fit

The RMSE of *Logi-ODE* on fitted years is quite low, although some local shape changes are missing. The averaged result for the given year and genotypes is RMSE scores of $0.045 \pm 1.13 \times 10^{-6}$ for the years 2018 and 2019, $0.021 \pm 4.33 \times 10^{-6}$ for the year 2022 and $0.033 \pm 2.55 \times 10^{-7}$ for the year 2021.

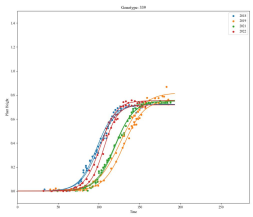

(a) genotype 339

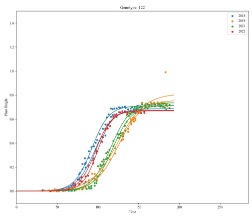

(b) genotype 122

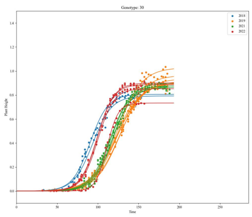

(c) genotype 30

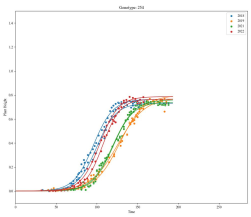

(d) genotype 254

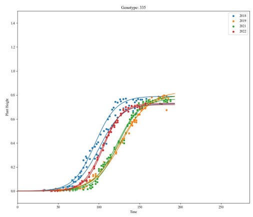

(e) genotype 335

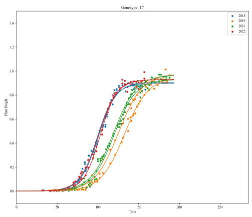

(f) genotype 17

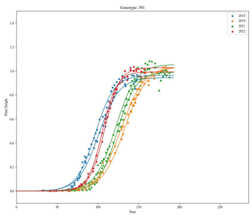

(g) genotype 301

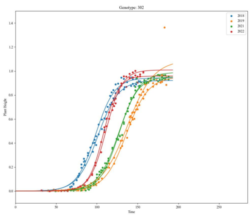

(h) genotype 302

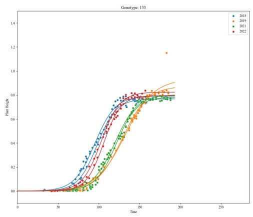

(i) genotype 133

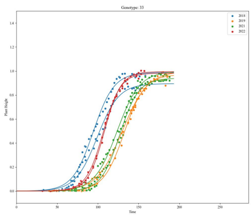

(j) genotype 33

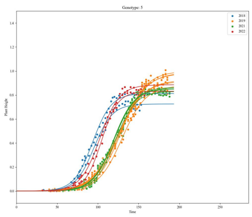

(k) genotype 5

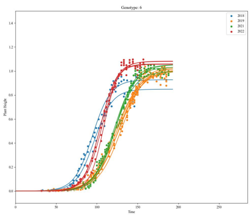

(l) genotype 6

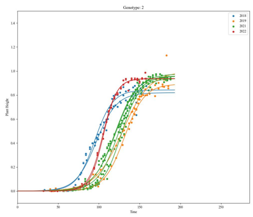

(m) genotype 2

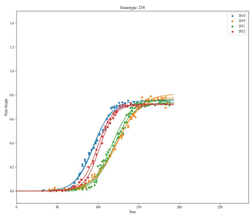

(n) genotype 218

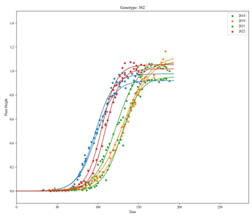

(o) genotype 362

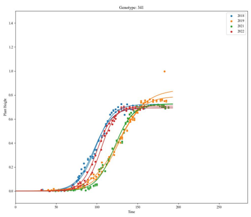

(p) genotype 341

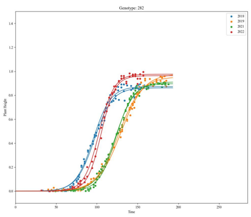

(q) genotype 282

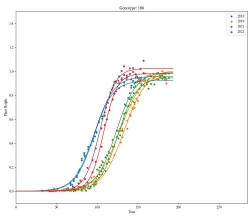

(r) genotype 106

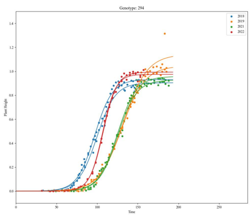

(s) genotype 294

**Figure S5.1:** *Logi-ODE* fit to each replicate (plot) separately for four years, which were ordered based on the same order as the result Figure 4. The dots are measured plant height data, and the curves are fitted with a logistic ODE to each sequence (plot/replicate) separately, coloured according to the harvest year. There is a clear difference between dates when the plants start to elongate and reach their maximum plant height. Unlike prediction models,we do not predict for new environmental conditions, but only want to justify that *Logi-ODE* can describe the general plant height growth curves.

#### S6. *Logi-PINN* model stability for different initialisation

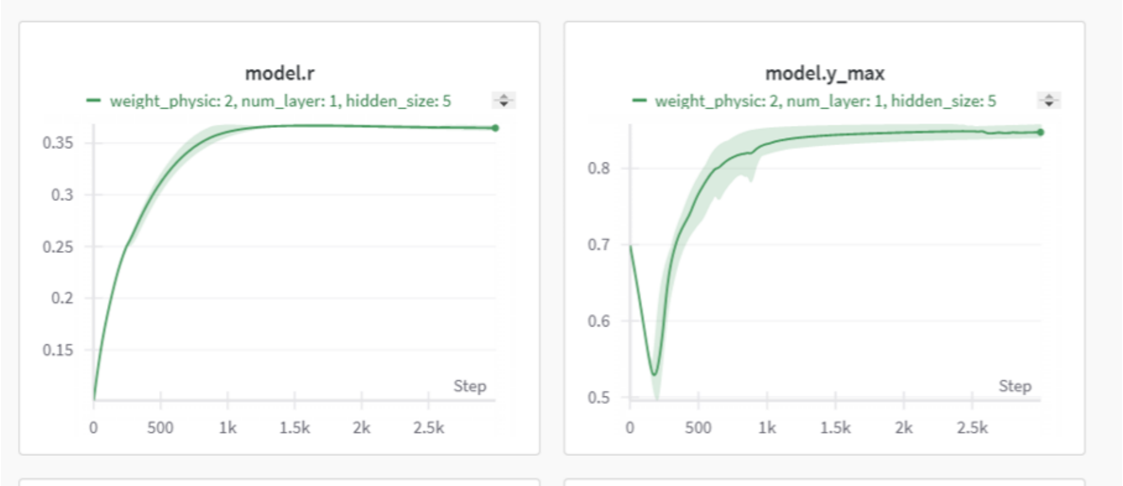


**Figure S6.1:** Parameter coverage for *Logi-PINN* model. This is an example from five runs with different random seeds of our *Logi-PINN*, which shows that $r(g)$ and $y_{max}(g)$ can converge to similar values for each genotype. The green line indicates the average $r(g)$ and $y_{max}(g)$ for a given genotype changing with the increase of training epochs, and the shade shows the actual value range (min and max).

#### S7. Weight parameters from trained *Logi-PINN* model for different genotypes

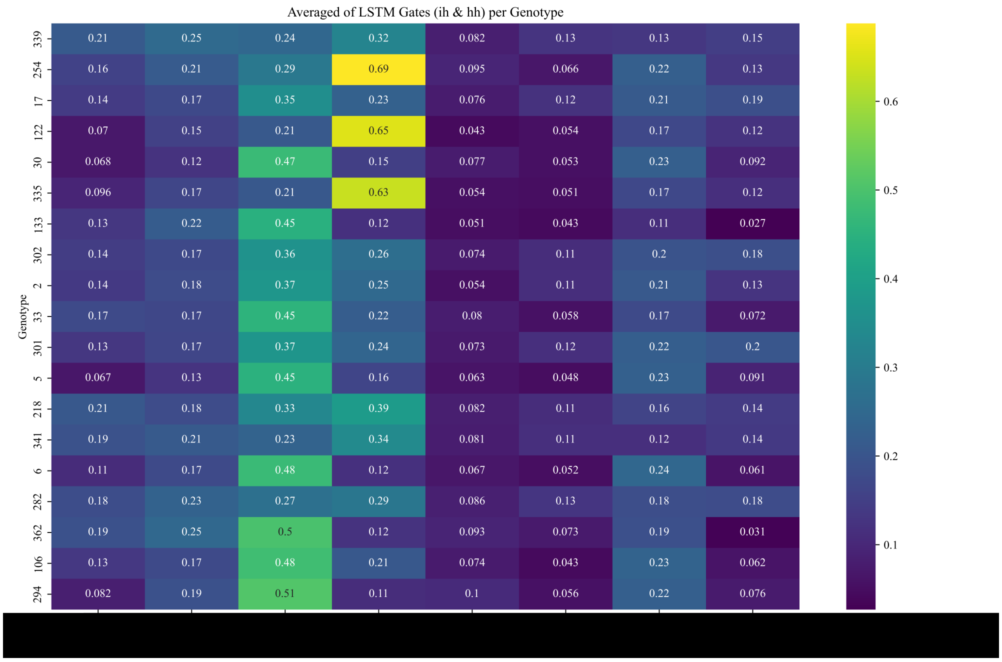


**Figure S7.1:** Heatmap of the averaged weights for the first LSTM layer from trained *Logi-PINN* models for different genotypes. The weight of the trained models implies how much information is used from previous time steps and the current time step. The weight values in the heatmap are normalised based on the weight values from the best hyperparameters and averaged across five models with different initialisations.

#### S8. Kinship matrix and genotype similarity

The kinship matrix we use represents genotype similarity based on the SNPs across their genome, which we also used to represent genetic effect and links to ODE parameters (for *Logi-PINN* structure) in multiple-genotype models. Those genotype effects are used to predict plant height development at different years. However, firstly, we found that the correlation (pairwise relatedness in kinship matrix) between the selected 19 genotypes is quite similar (all winter wheats), with no distinct groups within the 19 genotypes (Figure S8.1 (a)). Secondly, we also found no clear relationship between our calculated genotype similarity and *Logi-ODE* parameters (Figure S8.1 (b)).

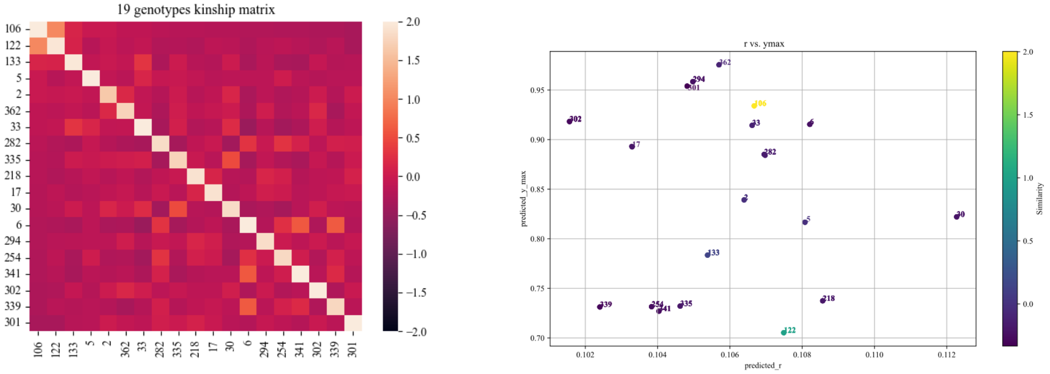


(a) kinship matrix

(b) ODE parameters from ODE solver

**Figure S8.1:** Kinship matrix and *Logi-ODE* parameters. The result section shows the results for the training years 2018 and 2019. The right heatmap does not show clear groups among different genotypes. There is also no clear relationship between ODE parameters and genotype similarity.
